# The origin and spread of locally adaptive seasonal camouflage in snowshoe hares

**DOI:** 10.1101/847616

**Authors:** Matthew R. Jones, L. Scott Mills, Jeffrey D. Jensen, Jeffrey M. Good

## Abstract

Adaptation is central to population persistence in the face of environmental change, yet we rarely precisely understand the origin and spread of adaptive variation in natural populations. Snowshoe hares (*Lepus americanus*) along the Pacific Northwest (PNW) coast have evolved brown winter camouflage through positive selection on recessive variation at the *Agouti* pigmentation gene introgressed from black-tailed jackrabbits (*L. californicus*). Here we combine new and published whole genome and exome sequences with targeted genotyping of *Agouti* in order to investigate the evolutionary history of local seasonal camouflage adaptation in the PNW. We find evidence of significantly elevated inbreeding and mutational load in coastal winter-brown hares, consistent with a recent range expansion into temperate coastal environments that incurred indirect fitness costs. The genome-wide distribution of introgression tract lengths supports a pulse of hybridization near the end of the last glacial maximum, which may have facilitated range expansion via introgression of winter-brown camouflage variation. However, signatures of a selective sweep at *Agouti* indicate a much more recent spread of winter-brown camouflage. Through simulations we show that the temporal lag between the hybrid origin and subsequent selective sweep of the recessive winter-brown allele can be largely attributed to the limits of natural selection imposed by simple allelic dominance. We argue that while hybridization during periods of environmental change may provide a critical reservoir of adaptive variation at range edges, the probability and pace of local adaptation will strongly depend on population demography and the genetic architecture of introgressed variation.

## Introduction

Local adaptation is fundamental to the persistence of populations during periods of rapid environmental change. In particular, local adaption to marginal habitats may increase a species’ niche breadth and range size (Holt and Gomulkiewicz 1997), enhancing their evolutionary resilience (Sgrò et al. 2011; Slatyer et al. 2013; Forsman 2016; Mills et al. 2018). Consequently, range edges where populations encounter marginal habitats and less favorable conditions may harbor crucial adaptive variation that facilitates long-term persistence in the face of environmental change (Hampe and Petit 2005; Hill et al. 2011; Cheng et al. 2014). Yet, range boundaries may also reflect the limits of natural selection if they are defined by environments where populations have failed to adapt (Antonovics 1976; Kirkpatrick and Barton 1997; Bridle and Vines 2007). Revealing how adaptive variation arises and spreads along range edges is, therefore, fundamental to understand the limitations of adaptation to new or changing environments (Ackerly 2003; Hampe and Petit 2005). However, we rarely possess detailed knowledge of the genetic basis and history of local adaptation in natural populations.

Several decades of theoretical research have established a framework for predicting demographic conditions along range margins, which are crucial in shaping population-level fitness and the potential for adaptation and range expansion. Populations inhabiting marginal habitats are generally predicted to be small and occur at low densities (Antonovics 1976; Kirkpatrick and Barton 1997), resulting in relatively reduced rates of beneficial mutation and levels of standing genetic variation (Pfennig et al. 2016). Small range-edge populations may further experience higher rates of inbreeding due to genetic drift (Wright 1931; Barton 2001) and accumulate deleterious variation (i.e., mutational load; Lynch et al. 1995; Willi et al. 2018), which can decrease the probability of population persistence (Mills and Smouse 1994). Elevated individual inbreeding and mutational load along range edges may also reflect past histories of adaptation and range expansion that result in non-equilibrium population dynamics. For instance, mating between close relatives may increase in founder populations that have recently undergone severe population contractions associated with range expansions (Frankham 1998). Likewise, mutational load may be amplified through the colonization of new environments because population contractions reduce the efficacy of selection against deleterious alleles at the expansion front (i.e., expansion load; Peischl et al. 2013; Henn et al. 2016; González-Martínez et al. 2017; Willi et al. 2018). Thus, when adaptation does occur along range margins it may produce negative feedbacks on population fitness and evolutionary potential.

Patterns of migration into range edge populations are also pivotal to their fitness and adaptive potential. Larger core populations are expected to produce relatively more migrants than smaller edge populations, leading to asymmetric rates of gene flow between core and peripheral habitats. In extreme scenarios, edge populations with low population growth rates (*λ*<1) can be demographic sinks that are maintained by immigration from the core of the range (Holt and Gomulkiewicz 1997; Griffin and Mills 2009). Highly asymmetric gene flow may further reduce fitness and hinder adaptation along the range edge by continually swamping local selection (Haldane 1930; Garcia-Ramos and Kirkpatrick 1997; Kirkpatrick and Barton 1997; Kawecki 2008). However, gene flow from core populations into edge populations may ultimately promote adaptive responses when edge populations are small and ecological gradients are shallow (Polechová 2018; Bontrager and Angert 2019). Hybridization between species may also facilitate adaptation and range expansion if edge populations intersect with the range of closely-related species that are adapted to local habitats (Baker 1948; Lewontin and Birch 1966; Burke and Arnold 2001; Rieseberg et al. 2007; Kawecki 2008; Pfennig et al. 2016). Introgression may provide a crucial source of large-effect variation (Hedrick 2013), which is predicted to be scarce in small populations but often necessary for range-edge adaptation and expansion (Behrman and Kirkpatrick 2011; Gilbert and Whitlock 2016). Putative adaptive introgression has now been shown in numerous species (e.g., Song et al. 2011; Pardo-Diaz et al. 2012; Huerta-Sánchez et al. 2014; Lamichhaney et al. 2015; Miao et al. 2016; Jones et al. 2018; Oziolor et al. 2019) and has been specifically linked to range expansions in Australian fruit flies (Lewontin and Birch 1966), sunflowers (Rieseberg et al. 2007), and mosquitoes (Besansky et al. 2003). While hybridization may facilitate adaptation and range expansion via large-effect mutations (Hedrick 2013; Nelson et al. 2019), the factors influencing the pace of adaptive introgression are often unclear.

Snowshoe hares (*Lepus americanus*) are broadly distributed across boreal and montane forests of North America. Most populations of hares undergo seasonal molts between brown (summer) and white (winter) coats to maintain crypsis in snow-covered environments. Seasonal camouflage is a crucial component of fitness in this system (Mills et al. 2013) as hares that become mismatched with their environment experience dramatically increased predation rates (i.e., 3-7% increase in weekly survival mortality; Zimova et al. 2016). However, some hares have adapted to mild winter environments by remaining brown in the winter (Mills et al. 2018). Brown winter camouflage in snowshoe hares is relatively rare across the entire range (<5% of the range), but predominant along portions of the southern range edge in the Pacific Northwest (PNW; Nagorsen 1983) with occurrence closely tracking regions of low seasonal snow cover (Mills et al. 2018). As snow cover across North America continues to decline under climate change, it is predicted that winter-brown camouflage may spread from the edge to the interior of the range, enhancing the evolutionary resilience of snowshoe hares (Jones et al. 2018; Mills et al. 2018). We previously demonstrated that brown versus white winter camouflage in PNW snowshoe hares is determined by a simple *cis*-regulatory polymorphism of the *Agouti* pigmentation gene that influences its expression during the autumn molt (Jones et al. 2018). The locally adaptive winter-brown allele is fully recessive and derived from introgressive hybridization with black-tailed jackrabbits (*Lepus californicus*), a closely-related scrub-grassland species that remains brown in the winter (Jones et al. 2018). Thus, the evolution of brown winter coats along coastal environments in the PNW represents one of the few verified cases of introgression underlying an adaptive trait of known ecological relevance in mammals (Taylor and Larson 2019).

The establishment of this genotype-to-phenotype link provides a powerful opportunity to examine how population history and hybridization shape local adaptation and expansion along the range edge. Here we seek to deepen our understanding of 1) the population history of PNW range edge snowshoe hares and 2) the origin and spread of winter-brown camouflage across coastal PNW environments. We first use previously published targeted exome data (61.7 Mb for 80 individuals; Jones et al. 2018) to estimate historical changes in population size (*N*), individual inbreeding coefficients, and mutational load in PNW hares. We then combine 11 whole genome sequences (WGS; six new and five previously published) with 61 newly assembled complete mitochondrial genomes and targeted genotyping of the introgressed *Agouti* region across 106 hares to resolve the timing of hybridization with black-tailed jackrabbits and the subsequent spread of winter-brown coat color variation. We use these data to test theoretical predictions for the maintenance and spread of adaptive variation in peripheral environments. Our study provides rare empirical insight into the dynamic interplay of environmental change, hybridization, and selection along range-edge environments and its evolutionary consequences.

## Methods

### Genomic data generation

All sample collection with live animals was performed under approved state permits and associated Animal Use Protocols approved through the University of Montana Institutional Animal Care and Use Committee (IACUC).

For some analyses, we used previously generated targeted whole exome data (61.7-Mb spanning 213,164 intervals; ∼25-Mb protein-coding exons, ∼28-Mb untranslated region, ∼9-Mb intronic or intergenic) for 80 snowshoe hares (21*×* mean coverage) collected from Washington (WA; *n*=13 winter-brown, *n*=13 winter-white), Oregon (OR; *n*=13 winter-brown, *n*=13 winter-white), Montana (MT; *n*=14 winter-white), and southwest British Columbia (BC; *n*=14 winter-brown; Jones et al. 2018). Hares from OR and WA were collected in the Cascade Range where populations are polymorphic for winter coat color (Fig. 1A). Hares from Seeley Lake in western MT are winter-white individuals, while those from BC were collected in low-lying regions near the Pacific coast where snowshoe hares are all winter-brown (Fig. 1A). To infer the history of hybridization, we performed whole genome resequencing of three black-tailed jackrabbits from OR and California (CA) and two winter-brown snowshoe hares from OR and BC. These samples complement WGS data previously generated for two black-tailed jackrabbits from Nevada (NV; one of which was sequenced to higher coverage in this study) and snowshoe hares from MT, WA, Utah (UT), and Pennsylvania (PA; Jones et al. 2018). We extracted genomic DNA following the Qiagen DNeasy Blood and Tissue kit protocol (Qiagen, Valencia, CA) and prepared genomic libraries following the KAPA Hyper prep kit manufacturer’s protocol. For all libraries, we sheared genomic DNA using a Covaris E220evolution ultrasonicator and performed a stringent size selection using a custom-prepared carboxyl-coated magnetic bead mix (Rohland and Reich 2012) to obtain average genomic fragment sizes of 400-500 bp. We determined indexing PCR cycle number for each library with quantitative PCR (qPCR) on a Stratagene Mx3000P thermocycler (Applied Biosystems) using a DyNAmo Flash SYBR Green qPCR kit (Thermo Fisher Scientific). Final libraries were size-selected again with carboxyl-coated magnetic beads, quantified with a Qubit (Thermo Fisher Scientific), and pooled for sequencing by Novogene (Novogene Corporation Ltd.; Davis, CA) on two lanes of Illumina HiSeq4000 using paired-end 150 bp reads.

**Figure 1.**
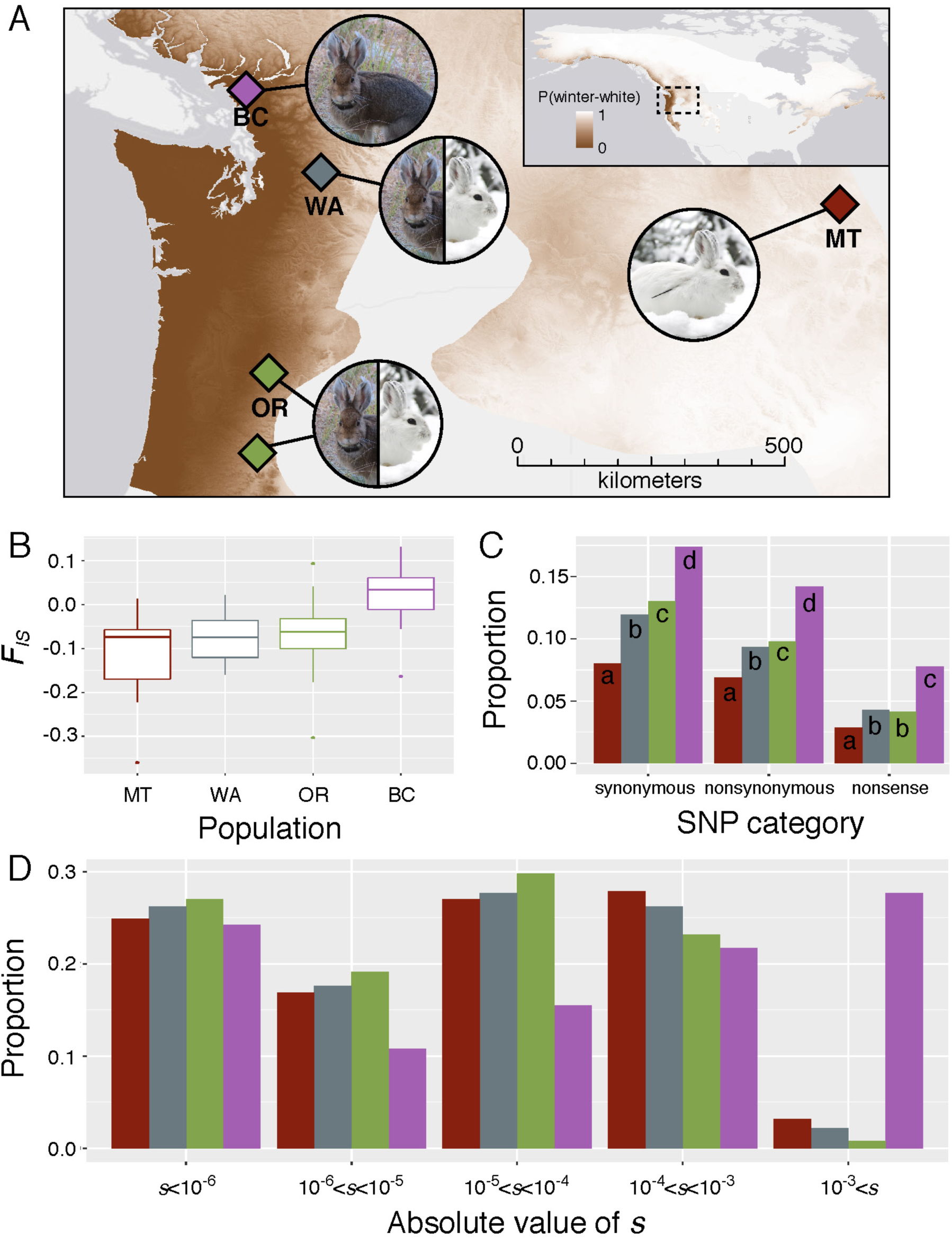
**(A)** Snowshoe hare range map colored by the probability of winter-white camouflage (white=1, brown=0). The Pacific Northwest region is magnified and shows sampling localities and coat color phenotypes for BC (purple), MT (red), OR (green), and WA (blue) populations used to generate whole exome data. (**B**) Box-and-whisker plots representing distributions of individual inbreeding coefficients (*F*_IS_) within each population. (**C**) The proportion of homozygosity across PNW populations for SNPs classified as synonymous, nonsynonymous (missense), or nonsense. Different letters denote significant differences between populations (*p*<0.01). (**D**) The inferred distribution of fitness effects for each population shown as the proportion of mutations with given selection coefficients.

To resolve the history of selection on the winter-brown *Agouti* allele, we performed targeted enrichment and sequencing to genotype 106 hares at the *Agouti* locus (*n*=37 WA, *n*=64 OR, *n*=5 MT). We developed a custom set of IDT xGen Lockdown probes spanning a 596.4 kb interval centered on the *Agouti* gene and extending to flanking regions (chromosome 4:5,250,800-5,847,200; coordinates based on the European rabbit (*Oryctolagus cuniculus*) oryCun2 genome build). The probe sequences were based on a snowshoe hare pseudoreference genome (∼33*×*; Jones et al. 2018) derived from iterative mapping to the rabbit genome (Carneiro et al. 2014). We targeted 96 uniquely-mapped 100 bp regions (based on low coverage WGS data from Jones et al. (2018)) that contained one or more diagnostic SNPs for winter coat color, allowing us to infer winter coat color for samples based on their *Agouti* genotype. We prepared genomic libraries for targeted *Agouti* sequencing following a modified version of Meyer and Kircher (2010), as described in Jones et al. (2018). We performed hybridization reactions on 500 ng of pooled libraries (10-16 individual libraries per pool), 5 μg of custom prepared snowshoe hare C_0_t-1 DNA, and 2 nM of blocking oligos. Washing and recovery of captured DNA was performed following the IDT xGen Lockdown probe hybridization capture protocol (version 2). Each capture library was then amplified in 50 μl reactions with 1X Herculase II reaction buffer, 250 μM each dNTP, 0.5 μM each primer, 1 μl Herculase II fusion polymerase, and 20 μl library template. The PCR temperature profile consisted of a 45 second 98°C denaturation step, followed by 24 cycles of 98°C for 15 seconds, 60°C for 30 seconds, and 72°C for 30 seconds, with a final 72°C elongation step for 1 minute. We cleaned and size-selected final libraries with 1.2X carboxyl-coated magnetic beads and verified target enrichment with qPCR. *Agouti* capture libraries were then pooled and sequenced with other libraries across two lanes of Illumina HiSeq4000 at the University of Oregon Core (Eugene, OR) and Novogene.

### Read processing and variant calling

For all raw sequence data, we trimmed adapters and low-quality bases (mean phred-scaled quality score <15 across 4 bp window) and removed reads shorter than 50 base pairs (bp) using Trimmomatic v0.35 (Bolger et al. 2014). We then merged paired-end reads overlapping more than 10 bp and with less than 10% mismatched bases using FLASH2 (Magoč and Salzberg 2011). Cleaned exome and *Agouti* capture reads were mapped using default settings in BWA-MEM v0.7.12 (Li 2013) to the snowshoe hare pseudoreference genome. WGS data were mapped to either the snowshoe hare or a black-tailed jackrabbit pseudoreference, which was also created by iteratively mapping to the rabbit genome (Jones et al. 2018). We used *PicardTools* to remove duplicate reads with the MarkDuplicates function and assigned read group information with the AddOrReplaceReadGroups function. Using GATK v3.4.046 (McKenna et al. 2010), we identified poorly aligned genomic regions with RealignerTargetCreator and performed local realignments with IndelRealigner. We performed population-level multi-sample variant calling using default settings with the GATK UnifiedGenotyper and filtered variants in VCFtools v0.1.14 (Danecek et al. 2011). For whole exome and whole genome data, we filtered genotypes with individual coverage <5*×* or >70*×* or with a phred-scaled quality score <30. Additionally, we removed all indel variants and filtered single nucleotide polymorphisms (SNPs) with a phred-scaled quality score <30 and Hardy-Weinberg *P*<0.001. We required that sites have no missing data across individuals. For targeted *Agouti* SNP data, we additionally filtered heterozygous genotypes with allelic depth ratios >3 and sites with > 50% missing data across individuals. We phased haplotypes and imputed missing data with Beagle v4.1 (Browning and Browning 2007) and used *Haplostrips* (Marnetto and Huerta-Sánchez 2017) to visualize haplotype structure.

### Population size history and inbreeding coefficients of PNW hares

We used the program *∂a∂i* (Gutenkunst et al. 2009) to infer historical population size (*N*) changes in PNW snowshoe hare populations (BC, MT, OR, and WA) using the folded site frequency spectrum (SFS) of synonymous variants from our extensive whole exome data set. We used the folded SFS to be consistent with statistical inferences of the distribution of fitness effects (see below). For each population, we tested a standard neutral equilibrium model, a two-epoch model (single instantaneous *N* change), a three-epoch model (two instantaneous *N* changes), an exponential *N* growth model, and an instantaneous *N* change + exponential *N* growth model. We inferred values for parameters*ν*, the population size relative to ancestral *N* (*N_anc_*; e.g., *ν*=1 if *N=N_anc_*) and *t*, the time of population size changes in units of 2*N_anc_* generations. We performed 100 independent runs under each model starting with parameter values sampled randomly across a uniform distribution (0.001<*ν*<100, 0<2*N_e_t*<2). For each model, we selected parameters with the highest log-likelihood value and chose the overall best model using a composite-likelihood ratio test with the Godambe Information Matrix (Coffman et al. 2016). We further checked the validity of maximum likelihood models by comparing the predicted SFS to the observed SFS for each population (Fig. S2). We determined 95% confidence intervals for parameter estimates using the Godambe Information Matrix with 100 bootstrap data sets comprised of one randomly selected synonymous SNP per 10 kb.

SFS-based approaches are often underpowered or inappropriate for inferring recent population size changes (Robinson et al. 2014; Beichman et al. 2018). For instance, even with a sufficient sample size, a historically large population that has very recently contracted in size (i.e., not in equilibrium) may nonetheless have a large variance *N_e_*. However, individuals in such populations may exhibit elevated individual inbreeding coefficients (*F*_IS_), calculated as 1-*H_o_*/*H_e_* where *H_o_* is the observed heterozygosity and *H_e_* is expected heterozygosity assuming random mating. To examine evidence for recent population contractions, we calculated the mean of the individual inbreeding coefficient (*F*_IS_) for each population using VCFtools and tested for significant differences between populations with a two-tailed Student’s t-Tests in R (R Core Team 2018).

### Mutational load and the distribution of fitness effects

For each PNW population, we measured the proportion of homozygosity across SNPs with predicted phenotypic effects (nonsynonymous and nonsense) as an indicator for relative differences in mutational load under a recessive deleterious mutation model (González-Martínez et al. 2017). We tested for significant differences in the proportion of homozygosity across populations using two-sided Z-tests for proportions in R (R Core Team 2018). Additionally, we used whole exome data to infer the distribution of selection coefficients of segregating variation, more commonly referred to as the distribution of fitness effects (DFE). In principle, we can infer the DFE from the SFS of selected sites because neutral, weakly deleterious, and strongly deleterious variation should segregate at different frequencies in populations (Keightley and Eyre-Walker 2010). The DFE of segregating variation is commonly inferred by first fitting a population history model to the SFS of neutral sites (often synonymous SNPs) and then fitting a mutational model to the SFS of selected sites (often nonsynonymous SNPs), while controlling for the effect of population history (i.e., changes in *N*) on the SFS of selected sites (Keightley and Eyre-Walker 2010). Here we implement this approach using the *Fit∂a∂i* module (Kim et al. 2017). We used the maximum likelihood parameter values from our inferred demographic model to control for population history and fit a simple DFE to the folded SFS of nonsynonymous variants (identified with SNPeff; Cingolani et al. 2012) described by a gamma distribution of selective effects with a shape (α) and scale (β) parameter. To estimate variance in shape and scale parameters, we used 100 bootstrap datasets randomly sampling 50% of nonsynonymous sites and performed 10 independent runs on each dataset. We used random starting values between 0.001 and 1 for the shape parameter and values between 0.01 and 200,000 for the scale parameter. To scale the DFE from relative selection coefficients (2*N_anc_s*) to absolute selection coefficients (*s*), we divided the scale parameter by 2*N_anc_* (Kim et al. 2017).

### The timing of hybridization

If hybridization between snowshoe hares and black-tailed jackrabbits is rare, then the age of hybridization may also reflect the age of *Agouti* introgression. We used two complementary approaches to estimate the timing of hybridization between PNW snowshoe hares and black-tailed jackrabbits. Previous phylogenetic analysis of partial cytochrome *b* sequences revealed that some PNW snowshoe hares carry introgressed mitochondrial DNA (mtDNA) genomes derived from hybridization with black-tailed jackrabbits (Cheng et al. 2014; Melo-Ferreira et al. 2014). We estimated the age of mtDNA introgression using complete mtDNA genomes for snowshoe hares (*n*=56) and black-tailed jackrabbits (*n*=5) that we assembled *de novo* from newly and previously generated WGS data (Jones et al. 2018) with the program NOVOPlasty (Dierckxsens et al. 2017). We aligned individual mtDNA assemblies, including the rabbit mtDNA reference as an outgroup (total assembled length= 16,251 bp), using default settings in Clustal W v2.1 (Larkin et al. 2007) and visually verified alignment quality. We then estimated a maximum clade credibility tree and node ages with a Calibrated Yule model in BEAST 2 (Bouckaert et al. 2014), assuming a strict molecular clock and an HKY substitution model using empirical base frequencies. We specified default priors for the kappa and gamma shape parameters and used a gamma distribution (alpha=0.001, beta=1000) as a prior for the clock rate and birth rate parameter. We ran the MCMC for 5 million steps and calibrated divergence times using a log-normal distribution for the rabbit-*Lepus* node age with a median of 11.8 million generations (95% prior density: 9.8–14.3; Matthee et al. 2004).

We also examined patterns of autosomal introgression tracts to infer the age of nuclear admixture. Given that mtDNA admixture may have been relatively ancient (Melo-Ferreira et al. 2014), admixture dating approaches based on linkage disequilibrium (LD) may have low power due to erosion of LD through ongoing recombination (Loh et al. 2013). Therefore, we developed an approach to fit the distribution of empirically inferred introgression tract lengths to tract lengths simulated under various models of admixture. We first identified genome-wide tracts of introgression using the program PhylonetHMM (Liu et al. 2014), which assigns one of two parent trees (species tree or hybridization tree) to each variable position using a hidden Markov model. PhylonetHMM robustly distinguishes between incomplete lineage sorting (ILS) and introgression by allowing for switches between gene trees within each parent tree (Liu et al. 2014; Schumer et al. 2016). Alignments of WGS data for the phylogenetic analysis included two black-tailed jackrabbits sampled from CA (BTJR1) and NV (BTJR2), a UT snowshoe hare (previously shown as non-admixed; Jones et al. 2018), and a winter-brown WA snowshoe hare to represent the admixed PNW snowshoe hare population. Here, the species tree is defined as ((WA,UT),(BTJR1, BTJR2)) and the hybridization tree is defined as (UT,(WA,BTJR1/BTJR2)). We specified base frequencies and transmission/transversion rates based on analysis with RAxML. We identified introgression tracts as contiguous regions of the genome with an average hybridization tree probability >95% across 25 variant windows (1 variant step) and excluded introgression tracts shorter than 10-kb (Schumer et al. 2016). We then used the program SELAM (Corbett-Detig and Jones 2016) to simulate a single pulse of admixture (lasting either 1 generation or 100 generations) occurring at a frequency of 0.01%, 0.1%, or 1% and recorded introgression tracts >10kb every 1000 generations for 50,000 generations across 21 autosomes. We performed a goodness of fit test between empirical and simulated tract length distributions using Kolmogorov-Smirnov tests (K-S tests), which measure differences in the cumulative fraction of data across the range of observed values (Massey 1951). To estimate the variance in hybridization timing, we performed K-S tests on 100 bootstrap data sets generated by subsampling 30% of the genome-wide introgression tracts.

### The time to most recent common ancestor of the winter-brown haplotype

To understand the history of the spread of winter brown camouflage, we used targeted *Agouti* SNPs to estimate the time to most recent common ancestor (TMRCA) for the winter-brown *Agouti* haplotype in OR (*n*=47 individuals) and WA (*n*=35 individuals). We estimated the TMRCA using a Markov chain Monte Carlo approach implemented in *startmrca* (Smith et al. 2018), which leverages information on the length distribution of the fixed selected haplotype and the accumulation of derived mutations. We assumed a constant recombination rate of 1 cM/Mb (Carneiro et al. 2011) and tested an upper and lower estimate for mutation rate in European rabbit (2.02 × 10^−9^ and 2.35 × 10^−9^ mutations/site/generation; Carneiro et al. 2012). We also explored the influence of using a divergent population (homozygous winter-white individuals from MT; *n*=5 individuals) or a local population (homozygous winter-white individuals from OR and WA; *n*=19 individuals) to represent the ancestral winter-white haplotype (Smith et al. 2018). We assumed chr4:5480355 (in oryCun2 coordinates) as the site of the “selected allele”, which lies in the center of the association interval between two strong candidate insertion-deletion mutations in the 5’ *cis*-regulatory region of *Agouti* and is perfectly correlated with winter coat color (Jones et al. 2018). We performed 100,000 iterations with a standard deviation of 20 for the proposal distribution and used the final 10,000 iterations to generate posterior TMRCA distributions (Smith et al. 2018).

### Simulations of selection on a recessive beneficial allele

Assuming fixation of a single haplotype, the above framework for inferring the TMRCA should reflect the age at which the beneficial haplotype began to increase rapidly in frequency (Smith et al. 2018), which under some conditions may be much more recent than the age of the beneficial mutation itself (Teshima and Przeworski 2006; Kelley 2012). For instance, the masking of recessive alleles to selection at low frequency is expected to decrease the rate at which they begin to increase in frequency, conditional on fixation (Teshima and Przeworski 2006), potentially resulting in a temporal lag between a fixed allele’s origin and TMRCA. However, such a scenario may be unlikely as the masking of rare recessive alleles is also expected to decrease their fixation probability (i.e., ‘Haldane’s sieve’, Haldane 1924; Turner 1981). Alternatively, an environmental change could favor a previously neutral or deleterious variant, resulting in a delayed spread of a segregating mutation. Indeed, Orr and Betancourt (2001) demonstrated that the bias against fixation of recessive alleles disappears when positive selection acts on pre-existing variation in mutation-selection balance. We used simulations to test whether different estimates of the timing of hybridization (i.e., the origin of the winter-brown haplotype) and TMRCA of the winter-brown allele could be due to the masking of recessive alleles at low frequency. Using SLiM 3.1 (Haller and Messer 2019), we simulated an equilibrium population (*N_e_*=257219 for OR population; Table 1) experiencing positive selection on a recessive allele (*s*=0.026, which reflects our updated median estimate of *s* for winter-brown haplotype in OR; Jones et al. 2018). At the beginning of the simulations, the recessive allele was introduced at a rate of 0.01% or 0.1% per generation for 1 or 100 generations, which reflects various rates and durations of hybridization. Under each hybridization scenario, we performed 100 simulations and tracked the frequency of the recessive allele every generation, conditioning on fixation. We saved tree sequences (Haller et al. 2019) and analyzed them using *msprime* (Kelleher et al. 2016) to identify the TMCRA for fixed beneficial alleles and determine whether selection resulted in fixation of a single copy (hard sweep) or multiple copies (soft sweep) of the beneficial allele.

**Table 1.**
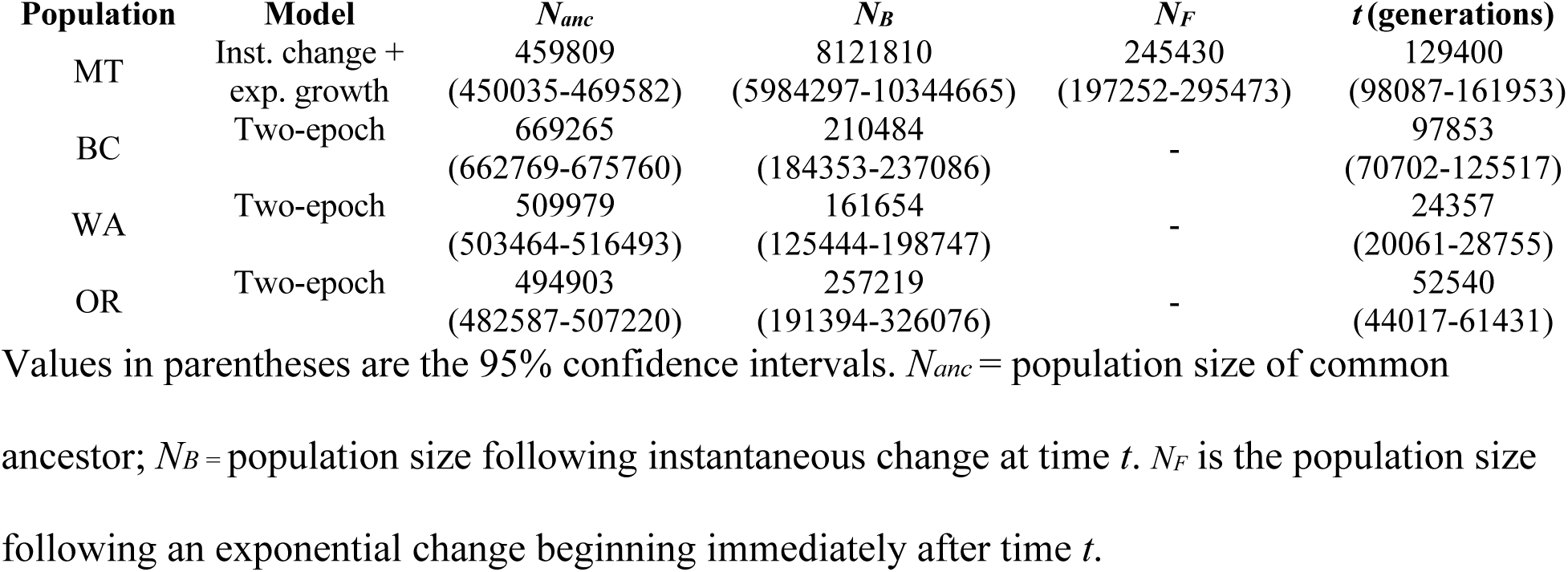
Maximum likelihood demographic model parameter estimates.

| Population | Model | $N_{anc}$ | $N_B$ | $N_F$ | $t$ (generations) |
| --- | --- | --- | --- | --- | --- |
| MT | Inst. change +<br>exp. growth | 459809<br>(450035-469582) | 8121810<br>(5984297-10344665) | 245430<br>(197252-295473) | 129400<br>(98087-161953) |
| BC | Two-epoch | 669265<br>(662769-675760) | 210484<br>(184353-237086) | - | 97853<br>(70702-125517) |
| WA | Two-epoch | 509979<br>(503464-516493) | 161654<br>(125444-198747) | - | 24357<br>(20061-28755) |
| OR | Two-epoch | 494903<br>(482587-507220) | 257219<br>(191394-326076) | - | 52540<br>(44017-61431) |

Values in parentheses are the 95% confidence intervals. $N_{anc}$ = population size of common ancestor; $N_B$ = population size following instantaneous change at time $t$ . $N_F$ is the population size following an exponential change beginning immediately after time $t$ .

## Results

### Range-edge population history and mutational load

We found support for a single relatively strong *N* contraction (i.e., a two-epoch model) occurring ∼24-100 kya in OR, WA, and BC hares (Table 1, Fig. S2). In contrast, the history of the MT population was characterized by an instantaneous + exponential *N* change model, in which the population experienced a sudden 17*×* expansion ∼129 kya followed by a gradual reduction to ∼53% of *N_anc_*. Despite population contractions, estimates of contemporary *N_e_* across all populations were relatively large (161654-257219; Table 1).

Using the same exome data set, we previously estimated the joint SFS for pairs of snowshoe hare populations to infer histories of population split times, migration rates, and effective population size in *∂a∂i* (Jones et al. 2018). These pairwise models supported histories of high symmetrical migration rates between populations and *N* contractions following population splits, but generated significantly smaller estimates of contemporary *N_e_* compared to the new estimates that we report in Table 1. However, we made a scaling error while estimating *θ* (= 4*N_e_μ*) under these previous models. This error affected our previously reported demographic parameter estimates for snowshoe hares (Table S9 in Jones et al. 2018) and associated selection coefficient parameter estimates (e.g., previous mean *s_WA_=*0.024, *s_OR_*=0.015; updated mean *s_WA_* =0.049, *s_OR_*=0.027; Durrett and Schweinsberg 2005; Jones et al. 2018), but not the main inference of introgression at *Agouti* underlying the genetic basis of polymorphic coat color in snowshoe hares. After scaling parameter values to the correct value of *θ* and excluding models beyond *a priori* divergence time parameter bounds (>500 thousand years), we found that our maximum likelihood demographic model (reported here in Table S1) still includes high migration rates between populations (∼1-2.63 migrants per generation), but with appreciably larger *N_e_* estimates than we previously reported and that are comparable to our new estimates (Table 1).

We found significantly elevated *F_IS_* in the coastal BC population compared to the other three PNW populations (*p*<0.01; Fig. 1B), which combined with our previous inference of elevated LD in this population (Jones et al. 2018) could suggest recent inbreeding and population size reduction. We further found a significantly higher proportion of homozygosity for synonymous, nonsynonymous, and nonsense SNPs in BC relative to other populations (Fig. 1C), which suggests elevated mutational load in BC under a recessive deleterious mutation model. BC individuals also have a significantly higher proportion of strongly deleterious nonsynonymous variants in (27.7%; |*s*|*≥* 10^−3^) relative to other populations (0.8-3.2%; Fig. 1D, Table S2). Because we have the same sample size for MT and BC (*n*=14 individuals) this striking difference in the DFE is likely not the result of the small BC sample size, which can lead to overestimation of the proportion of strongly deleterious variation (Kim et al. 2017). Notably, if synonymous SNPs used for demographic inference experience direct or linked selection (e.g., Akashi 1994; Stoletzki and Eyre-Walker 2006; Resch et al. 2007; Pouyet et al. 2018), then our demographic model could be misinferred (Ewing and Jensen 2016) in such a way that we will underestimate the strength of purifying selection on non-synonymous SNPs. Regardless, assuming levels of linked selection are similar across populations, the relative differences we observe in the DFE are unlikely driven by weak or linked selection on synonymous variants.

### The history of hybridization and introgression

From our complete mtDNA assemblies, we estimated a divergence time of 3.299 million generations ago (95% HPD interval: 2.555-4.255 million generations ago; Fig. 2) between black-tailed jackrabbit and non-introgressed snowshoe hares, which is consistent with previous estimates of species’ split times (Matthee et al. 2004; Melo-Ferreira et al. 2014; Jones et al. 2018). Within the non-introgressed snowshoe hare mtDNA clade, we found a relatively deep split between the UT snowshoe hare (representing the ‘Rockies’ cluster identified by Cheng et al. 2014) and all other snowshoe hares (641 thousand generations ago, 95% HPD interval: 476-834 thousand generations ago; Fig. 2).

**Figure 2.**
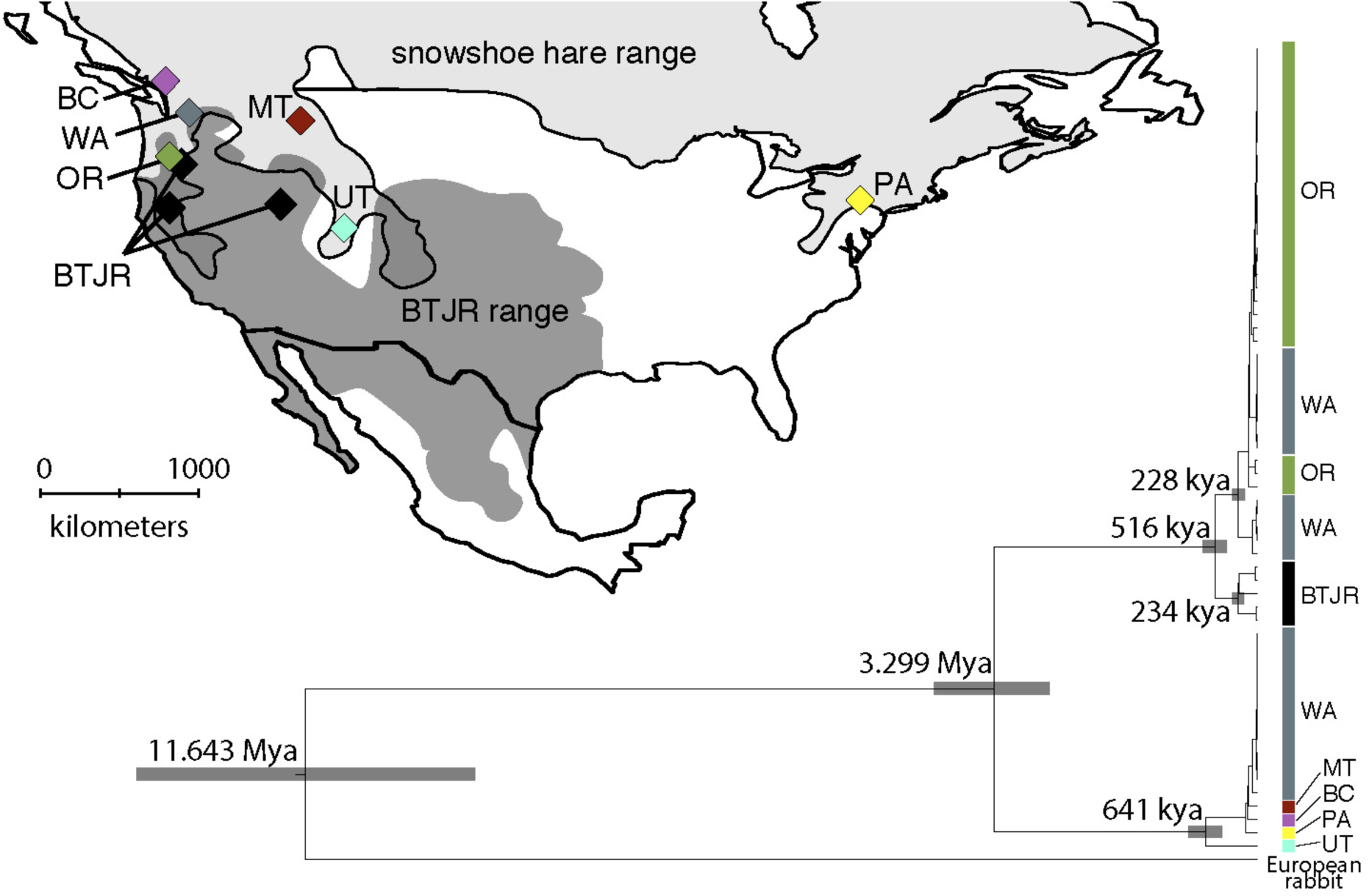
Snowshoe hare and black-tailed jackrabbit (BTJR) ranges with sampling localities for whole genome sequencing. The phylogenetic tree is a maximum clade credibility tree based on whole mitochondrial genome assemblies of snowshoe hares and black-tailed jackrabbits (European rabbit as outgroup) with median estimated split times for crucial nodes. Sample locality names and colors correspond to those on the map. Gray rectangles show the 95% highest posterior density (HPD) for each node age estimate.

A significant portion of snowshoe hares from the PNW (100% of OR hares and 50% of WA hares) formed a reciprocally monophyletic clade relative to black-tailed jackrabbits (100% posterior node support; Fig. 2). As previously demonstrated through coalescent simulations (Melo-Ferreira et al. 2014), this phylogenetic pattern cannot be plausibly explained by ILS and is consistent with asymmetric introgression of black-tailed jackrabbit mtDNA into snowshoe hares. As expected, mtDNA was not associated with winter coat color in the PNW polymorphic zone (chi-squared *P*=1). However, if we assume that hybridization is rare then mtDNA may track the same hybridization event that introduced winter-brown *Agouti* variation into PNW hares. The estimated split time between black-tailed jackrabbit and introgressed PNW hare mtDNA sequences was 516 thousand generations ago (95% HPD interval: 381-668 thousand generations ago, Fig. 2). However, this split time does not account for segregating ancestral polymorphism (Arbogast et al. 2002) or unsampled mtDNA variation within black-tailed jackrabbits. If we assume extant variation in snowshoe hares represents a single mtDNA introgression event, then the TMRCA of introgressed PNW snowshoe hare variation suggests a more recent date of mtDNA introgression of ∼228 thousand generations ago (95% HPD interval: 168-301 thousand generations ago).

Our previous work revealed elevated signatures of genome-wide nuclear admixture presumably coincident with introgression of seasonal camouflage variation (Jones et al. 2018). Here we identified 1878 individual introgression tracts (median length = 28,940 bp), encompassing ∼1.99% of the genome (Fig. 3). Across various simulated hybridization scenarios, the most strongly supported age of hybridization was 7-9 thousand generations ago with ranges of 95% confidence intervals spanning 6-10.5 thousand generations ago (Fig. S4). Different rates of admixture or admixture pulse lengths appeared to have little effect on the inferred hybridization age or the overall fit to empirical data (Fig. S4). Furthermore, we observed poor model fitting for very recent hybridization (< 5 thousand generations ago).

**Figure 3.**
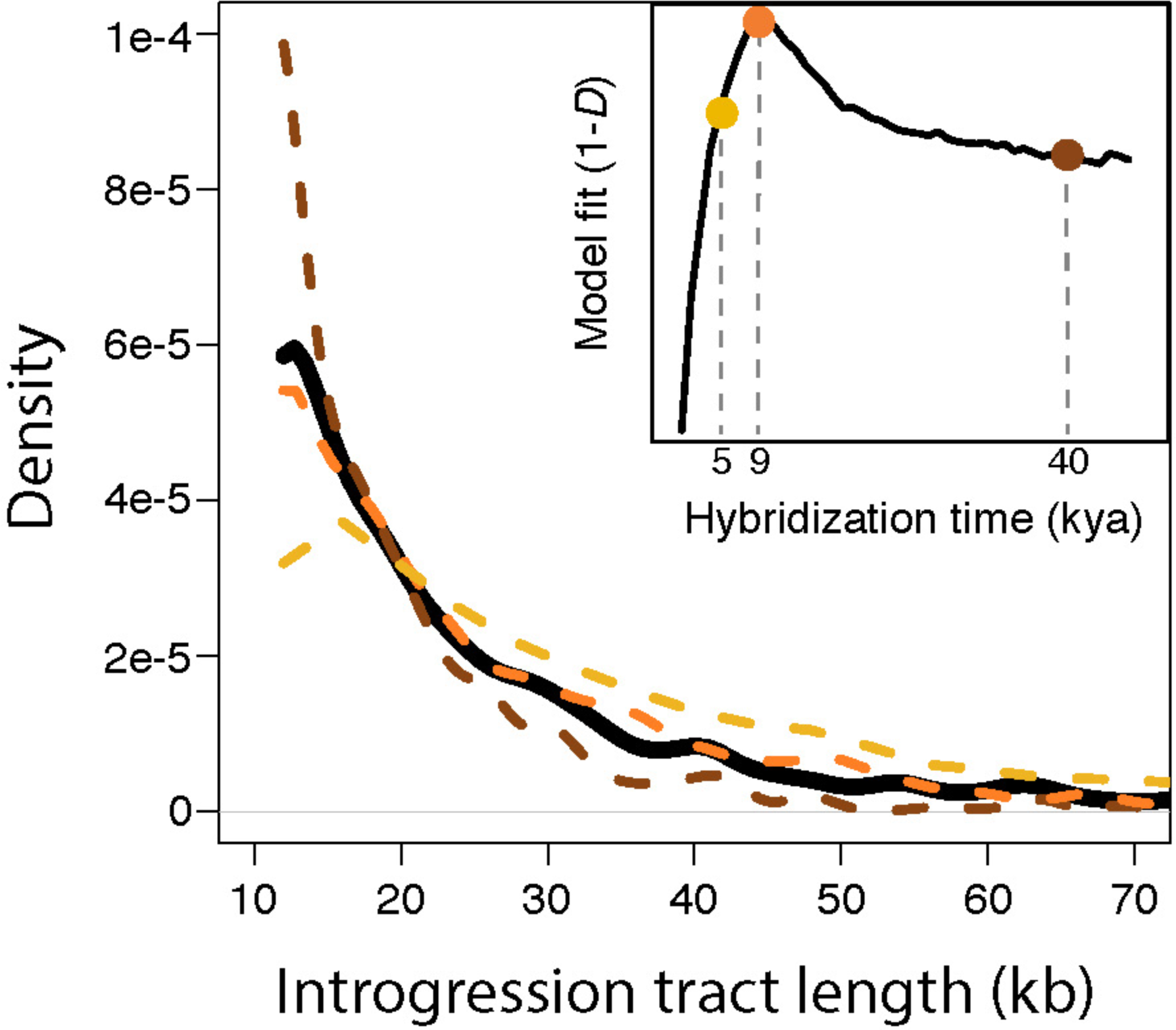
Empirical and simulated distributions of introgression tract lengths. The black line shows the empirical distribution of genome-wide introgression tract lengths in snowshoe hares. Colored dashed lines show simulated tract length distributions at three time points following a 100 generation pulse of hybridization at 0.1% frequency (yellow= 5 kya, orange=9 kya, brown=40 kya). The inset figure shows the overall fit of simulated introgression tract lengths to the empirical distribution through time, with hybridization occurring 9 kya as the model with the strongest fit (95% confidence intervals: 8-10.5 kya).

### Positive selection for winter-brown camouflage

We identified the *Agouti* region as one of the longest (209,012 bp) and most highly supported introgression tracts (mean introgression probability=0.99) in the WA winter-brown hare (Fig. 4). To understand the history of positive selection on brown winter camouflage, we estimated the TMRCA of the selected winter-brown *Agouti* haplotype in snowshoe hares using targeted sequencing across the *Agouti* region (mean coverage per interval 34*×* ± 17*×*). Using a divergent population, a local population, or both to represent the ancestral haplotype had little effect on TMRCA estimates (Table S3), so here we present estimates using both populations. Under a low or high estimate of the rabbit mutation rate, we inferred a TMRCA of approximately 1278 generations (95% CI: 1135-1441 generations) or 1226 generations (95% CI: 1054-1408 generations) for the winter-brown OR haplotype and approximately 1392 generations (95% CI: 1153-1607 generations) or 972 generations (95% CI: 766-1169 generations) for the WA haplotype, respectively (Table S3). We observed no consistent allelic differences between the fixed haplotypes in WA and OR (Fig. 4), consistent with a hard selective sweep.

**Figure 4.**
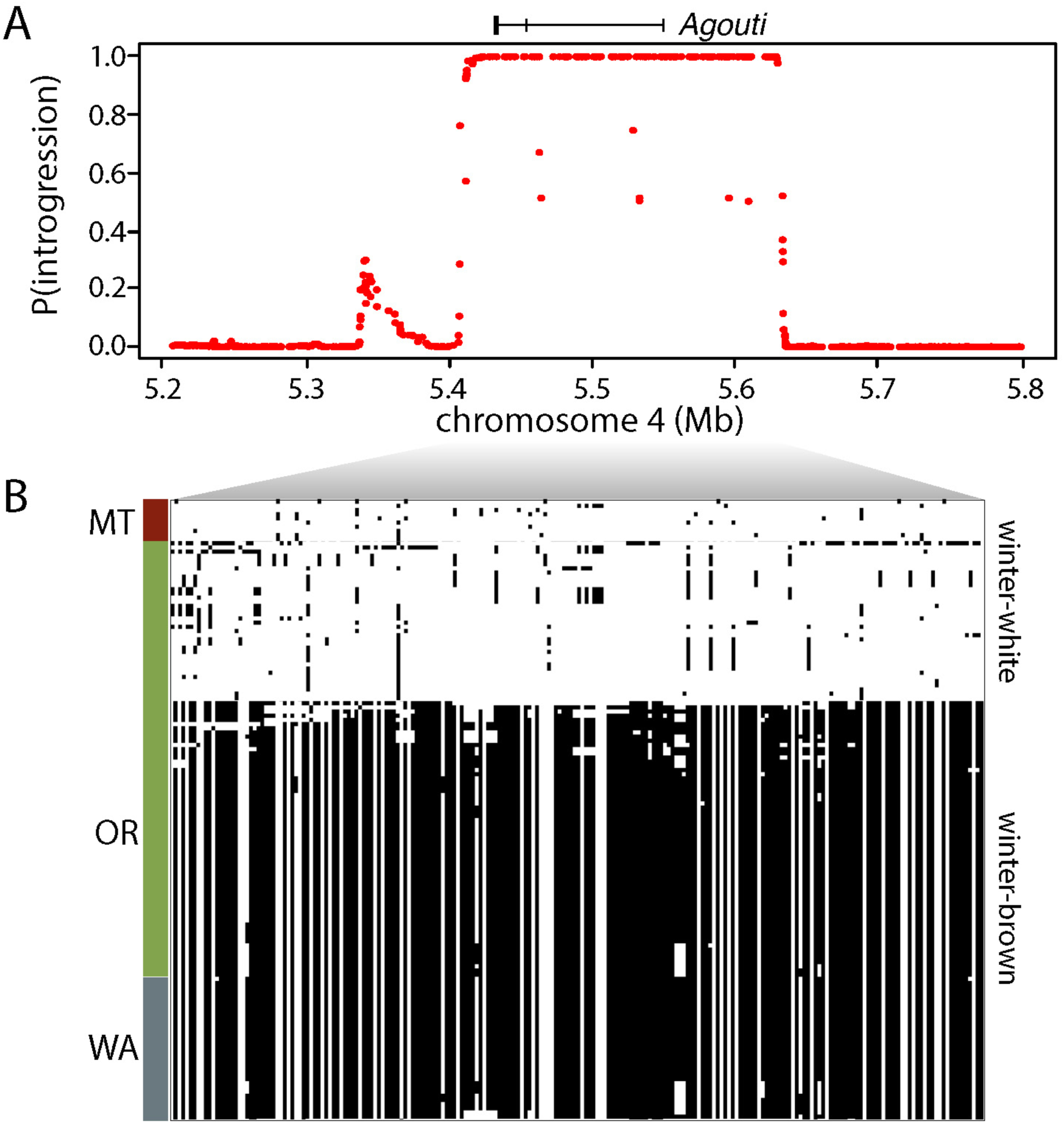
(**A**) Phylonet-HMM classification of the probability of introgression for each variable position across the *Agouti* locus. (**B**) Haplotype structure across the inferred introgressed *Agouti* interval (chr4:5424111-5633123) for winter-white and winter-brown PNW snowshoe hares.

Haplotype-based methods are known to underestimate the TMRCA and accounting for this systematic error produces TMRCA estimates of approximately 2-4 thousand generations (for a fully recessive allele, log_2_(estimate/true)*≈*−1.5; Kelley 2012) for the winter-brown haplotype in OR and WA. If our estimates are accurate, then there appears to be a ∼3-8 thousand generation lag between the origin of the winter-brown haplotype in snowshoe hares (i.e., the inferred hybridization date ∼ 7-9 thousand generations ago) and the increase in frequency of the winter-brown haplotype in the PNW from a single copy. Simulations show that such temporal lags are expected for selection on recessive variation, however the duration of this lag (and the total sojourn time) is negatively associated with the hybridization rate and fixation probability, as expected (Table 2). For instance, under the lowest hybridization rate (0.01% for 1 generation) the mean lag time was 2140 generations (95% CI: 101-8322 generation) with only a 0.8% fixation probability and under the highest hybridization rate (0.1% for 100 generations) the mean lag time was only 625 generations but with 100% fixation probability. Conditional on fixation, increased hybridization rates also tended to be more often associated with soft rather than hard sweeps (e.g., 38% hard sweeps for 0.1% hybridization rate for 100 generations versus 98% hard sweeps for 0.01% hybridization rate for 1 generation). However, under intermediate hybridization scenarios (0.1% for 1 generation or 0.01% for 100 generations), we observed relatively long mean lag times (2514 and 1587 generations, respectively) associated with high probabilities of fixation (12% and 79%, respectively), often through hard selective sweeps (81% and 96%; Table 2).

**Table 2.** Results from simulations of positive selection on recessive variation.

|  | **1 generation pulse** |  | **100 generation pulse** |  |
| --- | --- | --- | --- | --- |
|  | **0.01%** | **0.1%** | **0.01%** | **0.1%** |
| $P(\text{fixation})$ | 0.0082<br>(0.0068-0.010) | 0.12<br>(0.098-0.13) | 0.79<br>(0.71-0.85) | 1.0<br>(0.96-1.0) |
| $P(\text{hard sweep})$ | 0.98<br>(0.93-0.99) | 0.81<br>(0.72-0.87) | 0.96<br>(0.90-0.98) | 0.38<br>(0.29-0.48) |
| $T(\text{sojourn})$ | 8911<br>(5179-16208) | 9448<br>(5626-15746) | 6472<br>(5088-15339) | 5612<br>(4319-6757) |
| $T(\text{lag})$ | 2140<br>(101-8322) | 2514<br>(97-7740) | 1587<br>(87-7253) | 624<br>(91-1899) |

The beneficial variant was introduced through hybridization during a 1 or 100 generation pulse at a rate of 0.01% or 0.1%. Data are shown for the probability of fixation ( $P(\text{fixation})$ ), and, conditional on fixation, the probability of a hard selective sweep( $P(\text{hard sweep})$ ), the mean sojourn time ( $T(\text{sojourn})$ ), and the mean lag time between a fixed mutation's origin and TMRCA ( $T(\text{lag})$ ). In parentheses are 95% confidence intervals.

## Discussion

Range-edge adaptation may enhance a species’ evolutionary resilience to environmental change (Hampe and Petit 2005; Hill et al. 2011), however rigorous population genetic evaluations of predictions for range-edge demography and adaptation are limited (Bridle and Vines 2007). In snowshoe hares, the evolution of brown winter coats in temperate climates along the PNW coast represents the clearest example of local phenotypic adaptation in this wide-ranging species. Given its direct link to reduced snow cover, the evolution of brown winter camouflage may further foster persistence of snowshoe hares in the face of climate change (Mills et al. 2018). Here we leveraged our understanding of the genetic basis of brown winter camouflage to examine the history of range-edge adaptation, lending insights into the potential for rapid adaptation following environmental change.

### Population history and mutational load at the range edge

Populations along range margins are predicted to be small, limiting their ability to adapt to local conditions (although see Moeller et al. 2011; Graignic et al. 2018). Although we cannot assess relative differences in *N_e_* across the entire hare range, we uncovered high *N_e_* estimates across PNW populations (161654-257219; Table 1), despite evidence for strong ancient population size reductions. However, our *N_e_* estimates derive from predictions of genetic drift (i.e., variance *N_e_*) over long evolutionary time scales and may be a weak reflection of current census sizes, especially if local populations experience migration (Wang and Whitlock 2003) or have undergone recent size changes that are undetectable with the SFS (Beichman et al. 2018). We found evidence of significantly higher inbreeding coefficients and mutational load in coastal (BC) populations relative to the inland and montane populations (Fig. 1), signatures that are indicative of a recent population size reduction (Peischl et al. 2013, 2015; Bosshard et al. 2017; Gilbert et al. 2018). Elevated *F_IS_* and LD (Fig. S1 in Jones et al. 2018) could instead be related to cryptic population substructure (i.e., the Wahlund effect; Waples 2015). However, we have found no evidence for substructure or admixture in BC that could produce this effect (Jones et al. in prep). Similar signatures of elevated mutational load (e.g., homozygosity for deleterious alleles) have been found in other range-front populations, including the plant *Mercurialis annua* (González-Martínez et al. 2017) and in human populations that migrated out of Africa (Henn et al. 2016, although see Simons and Sella 2016). Thus, an intriguing potential explanation for these patterns is that they reflect signatures of a founder event associated with a recent range expansion. Moreover, given that we observe these signatures in the coastal winter-brown population, it is possible that this expansion was enabled by the evolution of locally adaptive brown winter camouflage. Winter-white hares experience heavy predation when mismatched (Zimova et al. 2016) and are not known to occur in low-lying coastal regional west of the Cascade Range (Nagorsen 1983; Mills et al. 2018), suggesting that coastal environments with ephemeral snow cover were likely unoccupied prior to local camouflage adaptation.

Long-term persistence of populations under environmental change ultimately requires adaptive evolution and the ability to colonize novel environments. If the colonization of coastal PNW environments by snowshoe hares was enabled by the evolution of brown winter coats, our results underscore that local adaptation to new environments can act as a negative feedback on fitness through the accumulation of deleterious mutations (Pujol and Pannell 2008; Gilbert et al. 2017; González-Martínez et al. 2017; Stewart et al. 2017; Willi et al. 2018). Although the consequences of mutational load for the persistence of PNW hare populations is unclear, high recessive mutational load may compromise the adaptive potential of populations (Assaf et al. 2015; González-Martínez et al. 2017) and increase the probability of extinction in small populations (Mills and Smouse 1994; Frankham 1998). In experiments of isolated *Tribolium* populations, short-term fitness gains via adaptive evolution were entirely lost over longer time periods as a consequence of increasing mutational load, although fitness could be readily restored through admixture (Stewart et al. 2017). In snowshoe hares, the potential fitness costs linked to mutational load may be mitigated by high gene flow between populations (Table S1) or superseded by the enhanced species-level evolutionary resilience afforded by brown-winter camouflage during periods of declining snow cover. Regardless, we suggest that any conservation efforts to promote adaptation to climate change should weigh the potential for enhanced long-term population persistence against the potential short-term fitness costs that may arise through mutational load.

### Hybridization and the origin of the winter-brown allele

Hybridization may play an important role shaping adaptation and expansion of range-edge populations (Pfennig et al. 2016), but evidence for this mode of adaptation stems from only a handful of examples (e.g., flies, Lewontin and Birch 1966; mosquitoes, Besansky et al. 2003; sunflowers, Rieseberg et al. 2007). In snowshoe hares, range and niche expansion into mild PNW coastal environments appears to have been enabled by adaptive introgression, although the history of hybridization has remained unclear. We estimated that mtDNA introgression in PNW snowshoe hares occurred ∼228 thousand generations ago, which could be interpreted as a conservative upper-bound for the timing of hybridization with black-tailed jackrabbits. Meanwhile, the genome-wide distribution of introgression tract lengths, which should be less sensitive to ILS and population structure within hares (Liu et al. 2014), suggest a much more recent pulse of hybridization ∼7-9 thousand generations ago (Fig. 3, Fig. S5). The different genome-wide and mtDNA estimates may also reflect independent pulses of ancient hybridization. Severe systematic overestimation of divergence dates may be common with mtDNA genomes calibrated with a relatively divergent outgroup because of high mutation rates and substitution saturation (Zheng et al. 2011). The divergence dates among major snowshoe hare mtDNA lineages also appear much deeper than our best estimates derived from population (nuclear) genomic data (∼2-3 fold deeper, unpublished data), which suggests that our analyses based on mtDNA likely overestimate the timing of introgression. We assumed a relatively simple molecular clock model and more complicated models (e.g., relaxed clocks) might better account for mutational processes observed in mtDNA genomes. However, it would seem that there would be little insight to be gained by additional modeling here given the myriad of limitations associated with extrapolating population history from a single stochastic realization of the coalescent process (Hudson and Turelli 2003).

Several recent studies have also noted that introgression is positively correlated with local recombination rate (Nachman and Payseur 2012; Janoušek et al. 2015; Schumer et al. 2018; Edelman et al. 2019; Li et al. 2019; Martin et al. 2019), presumable due to the effects of linked selection against deleterious mutations in hybrids. If this relationship generally holds, then it is possible that our dating approach based on the distribution of introgression tract lengths is also upwardly biased. However, contemporary range overlap between snowshoe hares and black-tailed jackrabbits appears restricted to relatively sharp ecological transitions between sage-scrub and montane forests in OR and CA (Fig. 2) and no records exist of putative hybrids, suggesting that contemporary hybridization is likely exceedingly rare or absent and has not resulted in discernable gene flow. Thus, the collective evidence suggests that historical hybridization between snowshoe hares and black-tailed jackrabbits was not more recent than 5 thousand generations ago. Notably, the timing of genome-wide admixture, assuming 1-2 generations per year in hares (Marboutin and Peroux 1995), appears coincident with the retreat of the Cordilleran ice sheet from low-lying coastal habitats in southern BC and northern WA at the end of the last glacial maximum (∼18 thousand years ago; Darvill et al. 2018) and thus the opening of suitable habitat for winter-brown snowshoe hares. This period of rapid climatic change resulted in individualistic range shifts for many North American mammal species (Graham 1986), potentially leading to novel community assemblages and thus promoting hybridization events (Swenson and Howard 2005), which could have created conditions favorable to adaptive introgression.

### The spread of winter-brown camouflage and the tempo of local adaptation

Although theory predicts adaptation in small range-edge populations may be slow and mutation-limited, hybridization may alleviate the lack of beneficial variation along range margins (Pfennig et al. 2016). Revealing how introgressed alleles adaptively spread through populations is therefore a critical component of understanding the limitations of range-edge adaptation. Here, we identified *Agouti* as one of the largest (>200 kb) and most strongly supported introgression tracts genome-wide (Fig. 4), consistent with our previous study showing exceptionally low genomic divergence in this region between black-tailed jackrabbits and winter-brown snowshoe hares (Jones et al. 2018). Assuming our genome-wide estimates of hybridization age reflect the origination of the *Agouti* allele through introgression (∼9 thousand generations ago), our findings suggest a ∼3-8 thousand generation delay until the selective sweep of the winter-brown haplotype in the PNW.

One potential biological explanation for this temporal lag is that winter-brown camouflage was not immediately beneficial in snowshoe hares. Rather, the winter-brown variant may have initially segregated as a neutral or deleterious allele for a period of time until an environmental shift allowed positive selection to act quickly on standing variation (e.g., Colosimo et al. 2005). However, our simulations suggest that beneficial recessive alleles segregating at frequencies as high as ∼10% (equivalent to simulations of 0.1% hybridization rate for 100 generations) take on average ∼5612 generations (95% CI: 4319-6757 generations) to reach fixation (Table 2). Thus, under an environmental shift scenario, the starting frequency of the winter-brown variant would likely have to be quite large (>10%) in order for selection to quickly drive it to fixation. Although allelic fixation under this model would be virtually guaranteed (Table 2), we suspect that such a high level of hybridization between black-tailed jackrabbits and snowshoe hares is unlikely given their ecological distinctiveness and our lower estimate of the genome-wide proportion of introgression (∼1.99%). Furthermore, the high starting allele frequency needed to result in rapid fixation is at odds with the evidence that selection fixed a single haplotype, as higher hybridization rates tended to result in softer sweeps (Table 2).

An alternative explanation for the delayed rise in frequency of the winter-brown allele invokes the limits of positive selection on recessive variation, which is predicted to result in an extended period of drift while at low frequency until homozygous recessive genotypes become more common. Consistent with this, we find significant temporal lags between the timing of hybridization and the TMRCA of fixed beneficial recessive alleles under low and moderate rates of hybridization. Although fixation under our lowest simulated rate of hybridization was highly unlikely (∼0.8%; Table 2), the two intermediate scenarios still resulted in relatively high fixation probabilities (12-78%) and tended to produce hard sweeps (81-96%), consistent with observed patterns of genetic variation at the winter-brown *Agouti* haplotype. These results demonstrate that one does not need to invoke changing selective coefficients to explain the apparent lag between the origin and the TMRCA of the winter-brown allele. Rather, our data are consistent with the winter-brown variant being immediately beneficial, although predominately hidden to selection, after introduced through hybridization at moderate frequency (∼0.1-1%). Indeed, this mutation-limited scenario is consistent with other known instances of colonization of novel environments through the evolution of locally adaptive camouflage in Nebraska deer mice and White Sands lizards (Laurent et al. 2016; Pfeifer et al. 2018; Harris et al. 2019).

Rates of adaptation at range edges are potentially an important component of species’ responses to climate change (Hampe and Petit 2005). Our study highlights the key role that hybridization can play in seeding adaptive variation and facilitating range expansion during periods of environmental change. In some cases, introgression appears to facilitate rapid adaptation to environmental change (Norris et al. 2015; Oziolor et al. 2019). However, introgression may not always be an efficient solution for rapid adaptation, as here we demonstrate that the rate of adaptation to novel mild winter environments in snowshoe hares appears to have been limited by the dominance coefficient of the winter-brown allele. Collectively, our findings demonstrate key factors that promote and limit adaptation to changing environments and, in particular, highlight the importance of characterizing genetic dominance of beneficial variants for understanding rates of adaptation and range expansion under climate change.

## Acknowledgements

We thank E. Cheng and K. Garrison for assistance with sample collection. We thank J. Melo-Ferreira, P. C. Alves, M. S. Ferreira, N. Herrera, E. Kopania, A. Kumar, M. Zimova, K. Garrison, N. Edelman, and the UNVEIL network for helpful discussions. We thank B. Kim for assistance with *Fit∂a∂i* analysis. Funding and support for this research was provided a National Science Foundation (NSF) Graduate Research Fellowship (DGE-1313190), NSF Doctoral Dissertation Improvement Grant (DGE-1702043), NSF Graduate Research Opportunities Worldwide, NSF EPSCoR (OIA-1736249), and NSF (DEB-0841884), the Drollinger-Dial Foundation, American Society of Mammalogists Grant-in-aid of Research, and a Swiss Government Excellence Scholarship. Original sequence data are available in the Sequence Read Archive (www.ncbi.nlm.nih.gov/sra). Previously generated whole exome and genome sequence data of snowshoe hare (BioProject PRJNA420081, SAMN02782769, SAMN07526959) are also available in the Sequence Read Archive.

