## Supplemental Figures/Tables for "The origin and spread of locally adaptive seasonal camouflage in snowshoe hares"

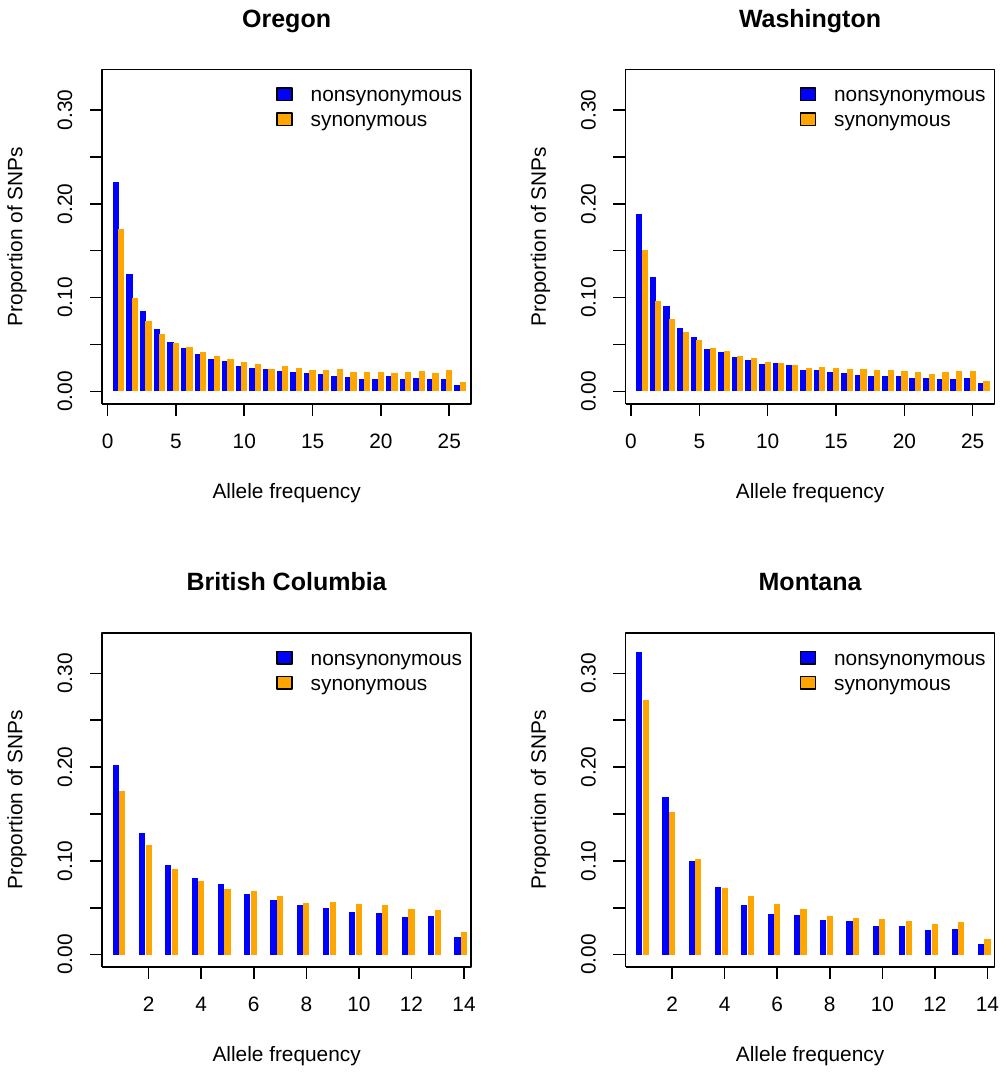

**Figure S1.** The site frequency spectrum for both synonymous (orange) and nonsynonymous (blue) variants for each PNW population. Synonymous variants were used to infer null demographic models while nonsynonymous variants were used to infer the distribution of fitness effects.

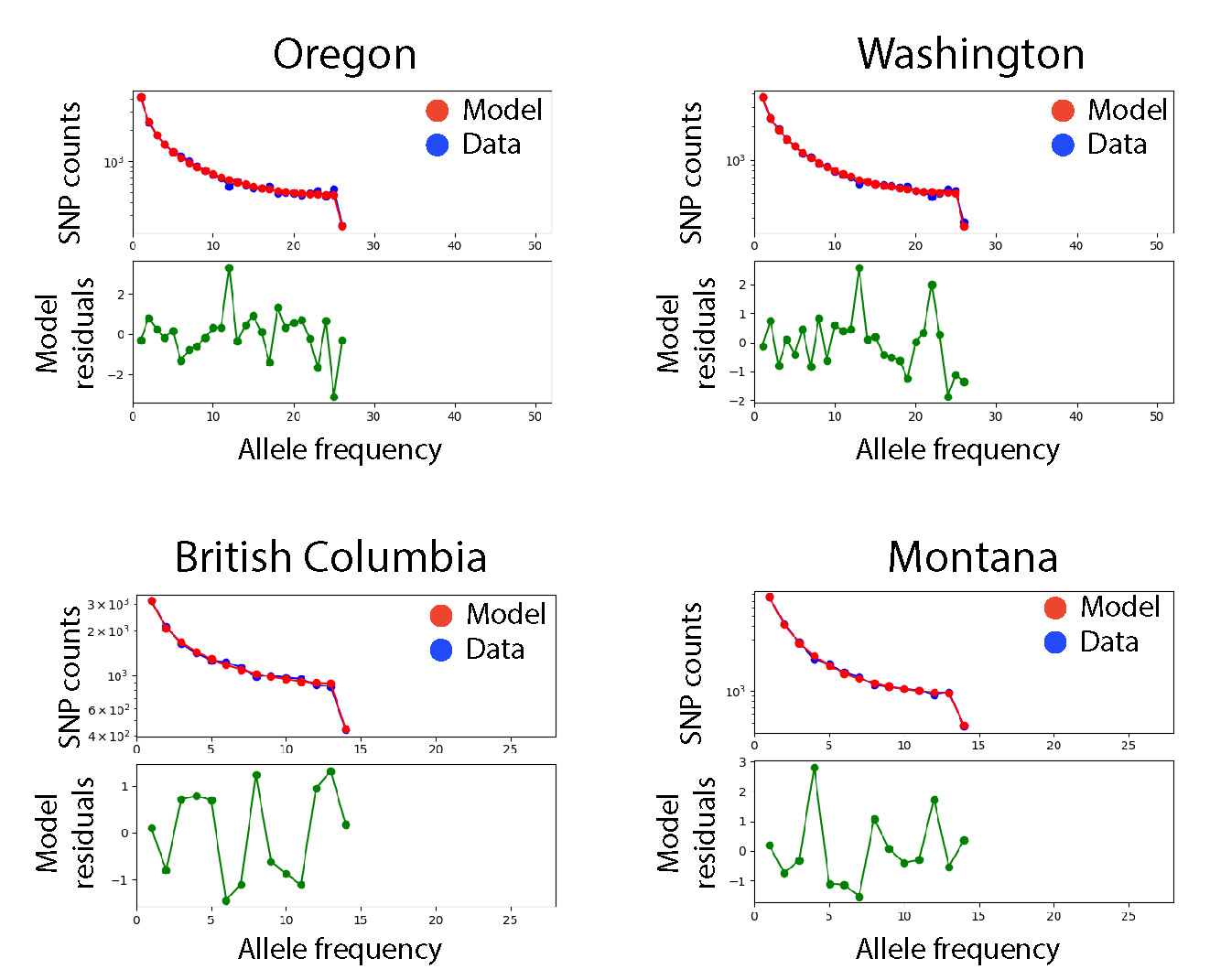

**Figure S2.** The fit of maximum likelihood demographic models to the empirical folded site frequency spectrum (SFS) for each population using synonymous SNPs. The predicted SFS under the demographic model is shown in red while the empirical SFS is shown in blue. Green points in plots below show the residuals.

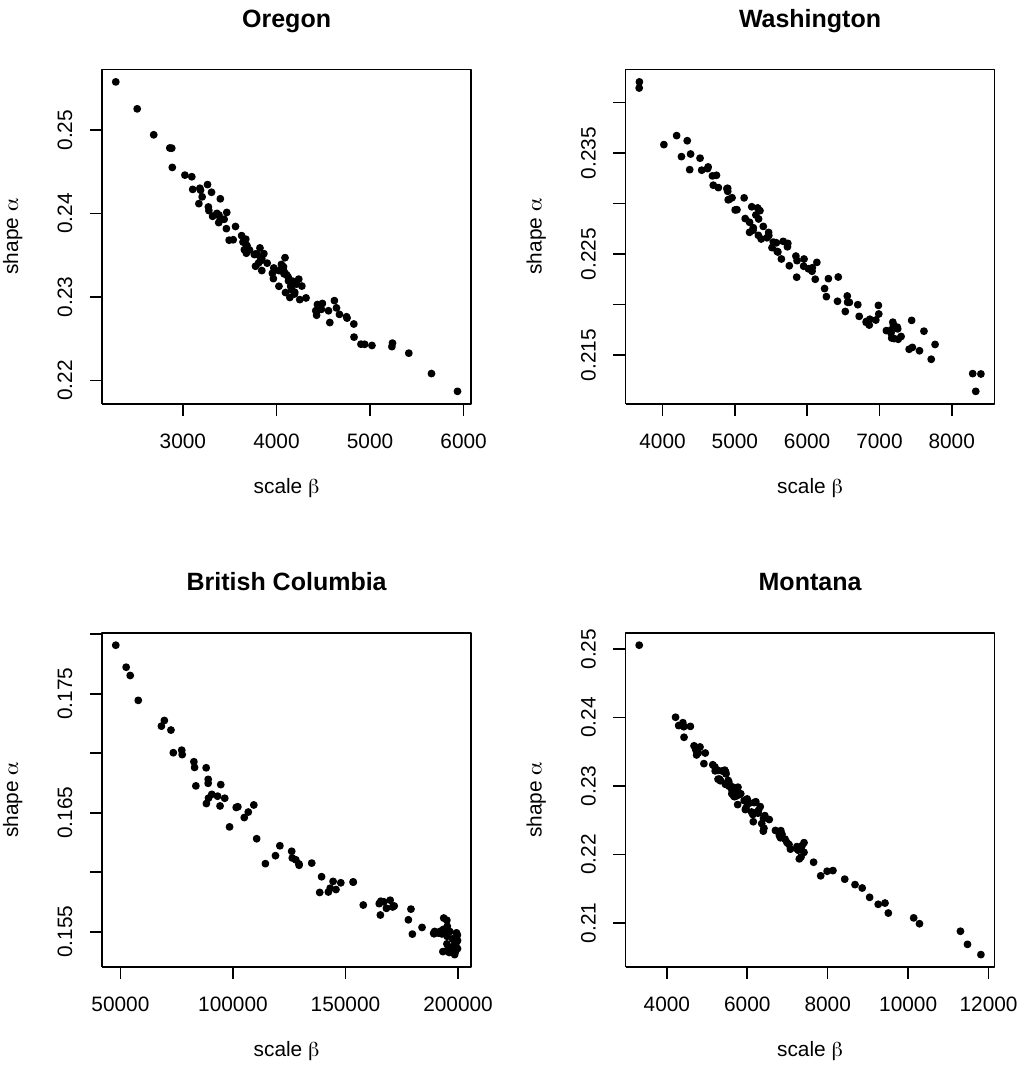

**Figure S3.** The estimated shape (α) and scale **(**β) parameters for gamma distributions describing the distribution of fitness effects for each population. Each point (100 per plot) shows a single estimate using a bootstrap data set of nonsynonymous variants.

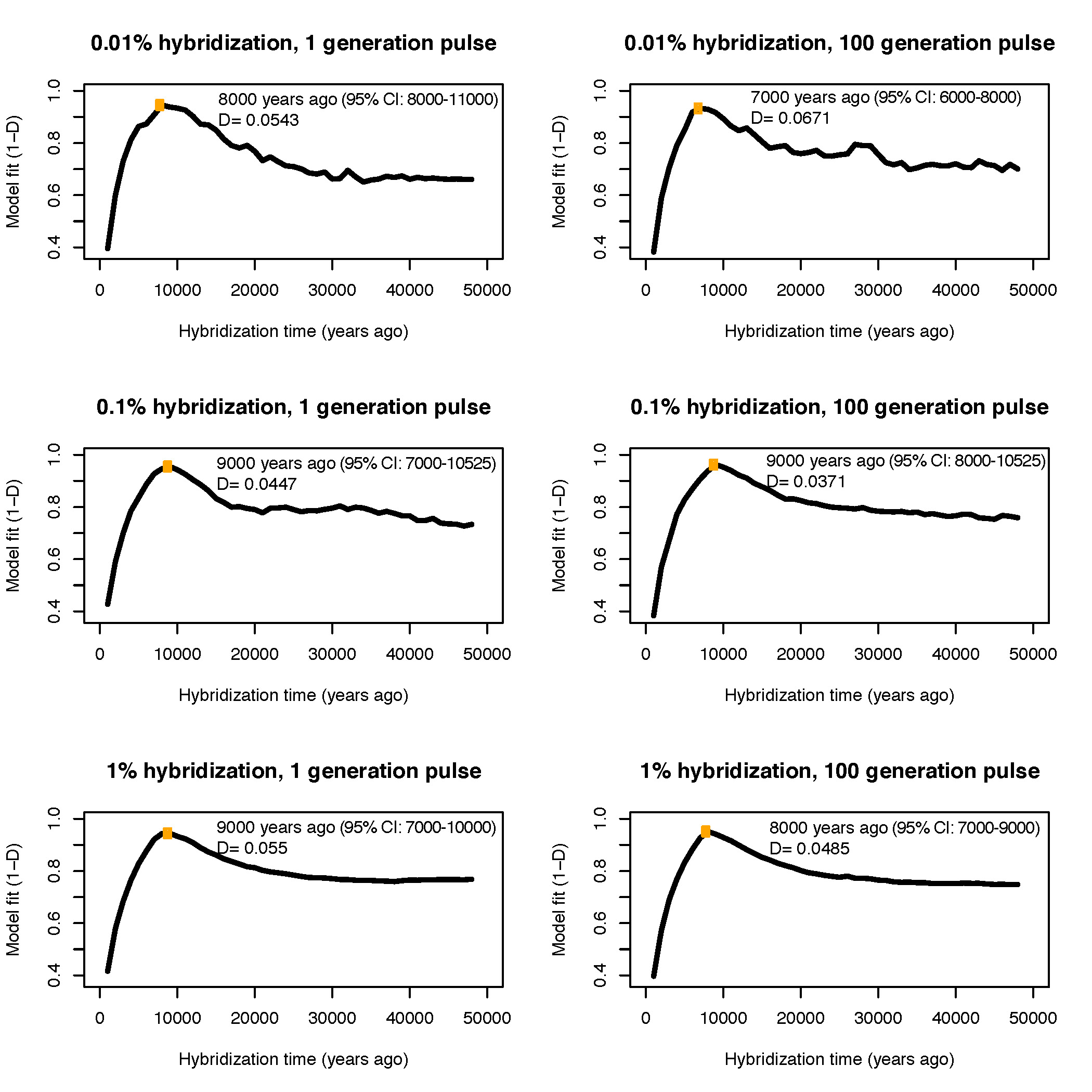

**Figure S4.** Introgression tract length model fits under either 0.01%, 0.1% or 1% hybridization rate lasting for 1 or 100 generations. Yellow dots indicate the optimal hybridization time in years (95% confidence intervals in parentheses) for each model, which is labelled in each figure along with the corresponding *D*-statistic from the Kolmogorov-Smirnov test. Lower values of *D* indicate better model fit.

**Table S1**. **Corrected maximum likelihood demographic model parameter estimates from Jones et al. (2018).** Values in parentheses represent 95% confidence intervals. *Nanc* = effective population size of common ancestor; *u_1_* = relative change in *Ne* of population 1 following split; *u_2_* = relative change in *Ne* of population 2 following split; *t* = divergence time in thousands of generations; *m* = the effective number of migrants per generation between populations.

| **P1-P2** | ***N*_anc_** | ***u*_1_** | ***u*_2_** | ***t*** | ***m*** |
| --- | --- | --- | --- | --- | --- |
| OR-WA | 304531  (301136-307925) | 0.23  (0.09-0.37) | 0.26  (0.11-0.40) | 30.89  (20.38-41.62) | 2.63  (0.33-4.94) |
| BC-WA | 288431  (284249-292613) | 0.10  (0-0.25) | 0.23  (0-0.63) | 15.26  (0-32.86) | 1.95  (0-8.12) |
| MT-WA | 339690  (337262-342119) | 0.52  (0.33-0.70) | 0.25  (0.10-0.41) | 54.56  (0.67-109.22) | 1.01  (0.39-1.63) |

**Table S2.** **Parameter estimates for the distribution of fitness effects.** The mean and 95% quantiles (in parentheses) are shown for the α and β parameters of the gamma distribution and the proportion of segregating variation binned by the absolute strength of selection (*s*).

| Population | α (shape) | β (scale) | 0≤\|*s*\|≤10^-6^ | 10^-6^≤\|*s*\|≤10^-5^ | 10^-5^≤\|*s*\|≤10^-4^ | 10^-4^≤\|*s*\|≤10^-3^ | 10^-3^≤\|*s*\| |
| --- | --- | --- | --- | --- | --- | --- | --- |
| MT | 0.226  (0.209-0.239) | 0.007  (0.004-0.012) | 0.249  (0.203-0.301) | 0.169  (0.051-0.283) | 0.270  (0.115-0.410) | 0.279  (0.157-0.387) | 0.032  (0.012-0.081) |
| BC | 0.160  (0.153-0.176) | 0.111  (0.042-0.149) | 0.242  (0.200-0.299) | 0.108  (0.001-0.225) | 0.155  (0.023-0.307) | 0.217  (0.063-0.387) | 0.277  (0.164-0.334) |
| WA | 0.225  (0.214-0.236) | 0.006  (0.004 - 0.008) | 0.262  (0.227-0.303) | 0.176  (0.087-0.267) | 0.277  (0.165-0.388) | 0.262  (0.178-0.333) | 0.022  (0.008-0.043) |
| OR | 0.234  (0.224-0.249) | 0.004  (0.003-0.005) | 0.270  (0.231-0.311) | 0.191  (0.097-0.286) | 0.298  (0.183-0.411) | 0.232  (0.160-0.298) | 0.008  (0.002-0.021) |

**Table S3.** Median TMRCA estimates for the winter-brown allele and 95% credible intervals using different mutation rates (2.02x10^-9^ or 2.35x10^-9^ mutations/base/generation) and homozygous winter-white samples to represent the ancestral reference (OR-WA as a ‘local’ reference and MT as a ‘divergent’ reference).

| Population | Reference | Mutation rate | TMRCA |
| --- | --- | --- | --- |
| OR | OR-WA | 2.02x10^-9^ | 1034 (920-1169) |
|  | OR-WA | 2.35x10^-9^ | 1038 (909-1176) |
|  | MT | 2.02x10^-9^ | 890 (785-1012) |
|  | MT | 2.35x10^-9^ | 1079 (958-1209) |
| WA | OR-WA | 2.02x10^-9^ | 1122 (945-1315) |
|  | OR-WA | 2.35x10^-9^ | 1101 (891-1300) |
|  | MT | 2.02x10^-9^ | 1067 (933-1221) |
|  | MT | 2.35x10^-9^ | 1354 (1174-1547) |
